# An approximation of the error back-propagation algorithm in a predictive coding network with local Hebbian synaptic plasticity

**DOI:** 10.1101/035451

**Authors:** James C.R. Whittington, Rafal Bogacz

## Abstract

To efficiently learn from feedback, cortical networks need to update synaptic weights on multiple levels of cortical hierarchy. An effective and well-known algorithm for computing such changes in synaptic weights is the error back-propagation algorithm. However, in the back-propagation algorithm, the change in synaptic weights is a complex function of weights and activities of neurons not directly connected with the synapse being modified, whereas the changes in biological synapses are determined only by the activity of pre-synaptic and post-synaptic neurons. Several models have been proposed that approximate the back-propagation algorithm with local synaptic plasticity, but these models require complex external control over the network or relatively complex plasticity rules. Here we show that a network developed in the predictive coding framework can efficiently perform supervised learning fully autonomously, employing only simple local Hebbian plasticity. Furthermore, for certain parameters, the weight change in the predictive coding model converges to that of the back-propagation algorithm. This suggests that it is possible for cortical networks with simple Hebbian synaptic plasticity to implement efficient learning algorithms in which synapses in areas on multiple levels of hierarchy are modified to minimize the error on the output.

## Introduction

Efficiently learning from feedback often requires changes in synaptic weights in many cortical areas. For example, when a child learns sounds associated with letters, after receiving a feedback from a parent, the synaptic weights need to be modified not only in auditory areas, but also in associative and visual areas. An effective algorithm for supervised learning of desired associations between inputs and outputs in networks with hierarchical organization is the error back-propagation algorithm (Rumelhart et al., 1986). Artificial neural networks (ANNs) employing back-propagation have been used extensively in machine learning (LeCun et al., 1989; Chauvin and Rumelhart, 1995; Bogacz et al., 1999), and have become particularly popular recently, with the newer deep networks having some spectacular results, now able to equal and out-perform humans in many tasks (Krizhevsky et al., 2012; Hinton et al., 2012). Furthermore, models employing the back-propagation algorithm have been successfully used to describe learning in the real brain during various cognitive tasks (Seidenberg and McClelland, 1989; McClelland et al., 1995; Plaut et al., 1996).

However, it has not been known if natural neural networks could employ an algorithm analogous to the back-propagation used in ANNs. In ANNs, the change in each synaptic weight during learning is calculated by a computer as a complex, global function of activities and weights of many neurons (often not connected with the synapse being modified). In the brain however, the network must perform its learning algorithm locally, on its own without external influence, and the change in each synaptic weight must depend just on the activity of pre-synaptic and post-synaptic neurons. This led to a common view of the biological implausibility of this algorithm (Crick, 1989), e.g. “despite the apparent simplicity and elegance of the back-propagation learning rule, it seems quite implausible that something like equations […] are computed in the cortex” (p. 162) (O’Reilly and Munakata, 2000).

Several researchers aimed at developing biologically plausible algorithms for supervised learning in multilayer neural networks. However, the biological plausibility was understood in different ways by different researchers, thus to help us evaluate the existing models, we define the criteria we wish a learning model to satisfy, and we will consider the existing models within these criteria.

1. Local computation: A neuron only performs computation on the basis of the inputs it receives from other neurons weighted by the strengths of its synaptic connections.
2. Local plasticity: The amount of synaptic weight modification is only dependent on the activity of the two neurons the synapse connects (and possibly a neuromodulator).
3. Minimal external control: The neurons perform the computation autonomously with as little external control routing information in different ways at different times as possible.
4. Plausible architecture: The connectivity patterns in the model should be consistent with basic constraints of connectivity in neocortex.

The models proposed for supervised learning in biological multilayer neural networks can be divided in two classes. Models in the first class assume that neurons (Barto and Jordan, 1987; Mazzoni et al., 1991; Williams, 1992) or synapses (Unnikrishnan and Venugopal, 1994; Seung, 2003) behave stochastically, and receive a global signal describing the error on the output (e.g. via a neuromodulator). If the error is reduced, then the weights are modified to make the produced activity more likely. Many of these models satisfy the above criteria, but they do not directly approximate the back-propagation algorithm, and it has been pointed that under certain conditions their learning is slow and scales poorly with network size (Werfel et al., 2005). The models in the second class explicitly approximate the back-propagation algorithm (O’Reilly, 1998; Lillicrap et al., 2014; Balduzzi et al., 2014; Bengio, 2014; Bengio et al., 2015; Scellier and Bengio, 2016), and we will compare them in detail in the Discussion.

Here we show how the back-propagation algorithm can be closely approximated in a model that uses a simple local Hebbian plasticity rule. The model we propose is inspired by the predictive coding framework (Rao and Ballard, 1999; Friston, 2003, 2005). The predictive coding framework is related to the auto-encoder framework (Ackley et al., 1985; Hinton and McClelland, 1988; Dayan et al., 1995) in which the GeneRec (O’Reilly, 1998) and another approximation of back-propagation (Bengio, 2014; Bengio et al., 2015) were developed. In both frameworks the networks include feed-forward and feedback connections between nodes on different levels of hierarchy, and learn to predict activity on lower levels from the representation on the higher levels. The predictive coding framework describes a network architecture in which such learning has a particularly simple neural implementation. The distinguishing feature of the predictive coding models is that they include additional nodes encoding the difference between the activity on a given level and that predicted by the higher level, and that these prediction errors are propagated through the network (Rao and Ballard, 1999; Friston, 2005). Patterns of neural activity similar to such prediction errors have been observed during perceptual decision tasks (Summerfield et al., 2006, 2008). In this paper we show that when the predictive coding model is used for supervised learning, the prediction error nodes have activity very similar to the error terms in the back-propagation algorithm. Therefore, the weight changes required by the back-propagation algorithm can be closely approximated with simple Hebbian plasticity of connections in the predictive coding networks.

In the next section we review back-propagation in ANNs. Then we describe a network inspired by the predictive coding model, in which the weight update rules approximate those of conventional back-propagation. We point out that for certain architectures and parameters, learning in the proposed model converges to the back-propagation algorithm. We compare the performance of the proposed model and the ANN. Furthermore, we characterize the performance of the predictive coding model in supervised learning for other architectures and parameters, and highlight that it allows learning bidirectional associations between inputs and outputs. Finally, we discuss the relationship of this model to previous work.

## Models

While we introduce ANNs and predictive coding below, we use a slightly different notation than in their original description to highlight the correspondence between the variables in the two models. The notation will be introduced in detail as the models are described, but for reference it is summarized in Table 1. To make dimensionality of variables explicit we denote vectors with a bar (e.g. ). MATLAB codes simulating an ANN and the predictive coding network are freely available at the ModelDB repository with access code 218084.

**Table 1:** Corresponding and common symbols used in describing ANNs and predictive coding models.

|  | Back-propagation | Predictive coding |
| --- | --- | --- |
| Activity of a node (before non-linearity) | $y_i^{(l)}$ | $x_i^{(l)}$ |
| Synaptic weight | $w_{i,j}^{(l)}$ | $\theta_{i,j}^{(l)}$ |
| Objective function | $E$ | $F$ |
| Prediction error | $\delta_i^{(l)}$ | $\varepsilon_i^{(l)}$ |
| Activation function | $f$ | |
| Number of neurons in a layer | $n^{(l)}$ | |
| Highest index of a layer | $l_{max}$ | |
| Input from the training set | $s_i^{in}$ | |
| Output from the training set | $s_i^{out}$ | |

## Review of error back-propagation

ANNs (Rumelhart et al., 1986) are configured in layers, with multiple neuron-like nodes in each layer as illustrated in Figure 1A. Each node gets input from a previous layer weighted by the strengths of synaptic connection, and performs a non-linear transformation of this input. To make link with predictive coding more visible, we change the direction in which layers are numbered, and index the output layer by 0 and the input layer by *l_max_*. We denote by 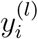 the input to the *i^th^* node in the *l^th^* layer, while the transformation of this by an activation function is the output, 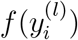. Thus:

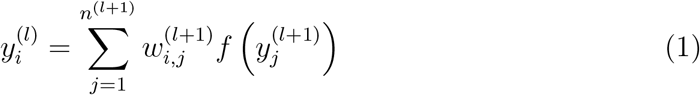

**Figure 1:**
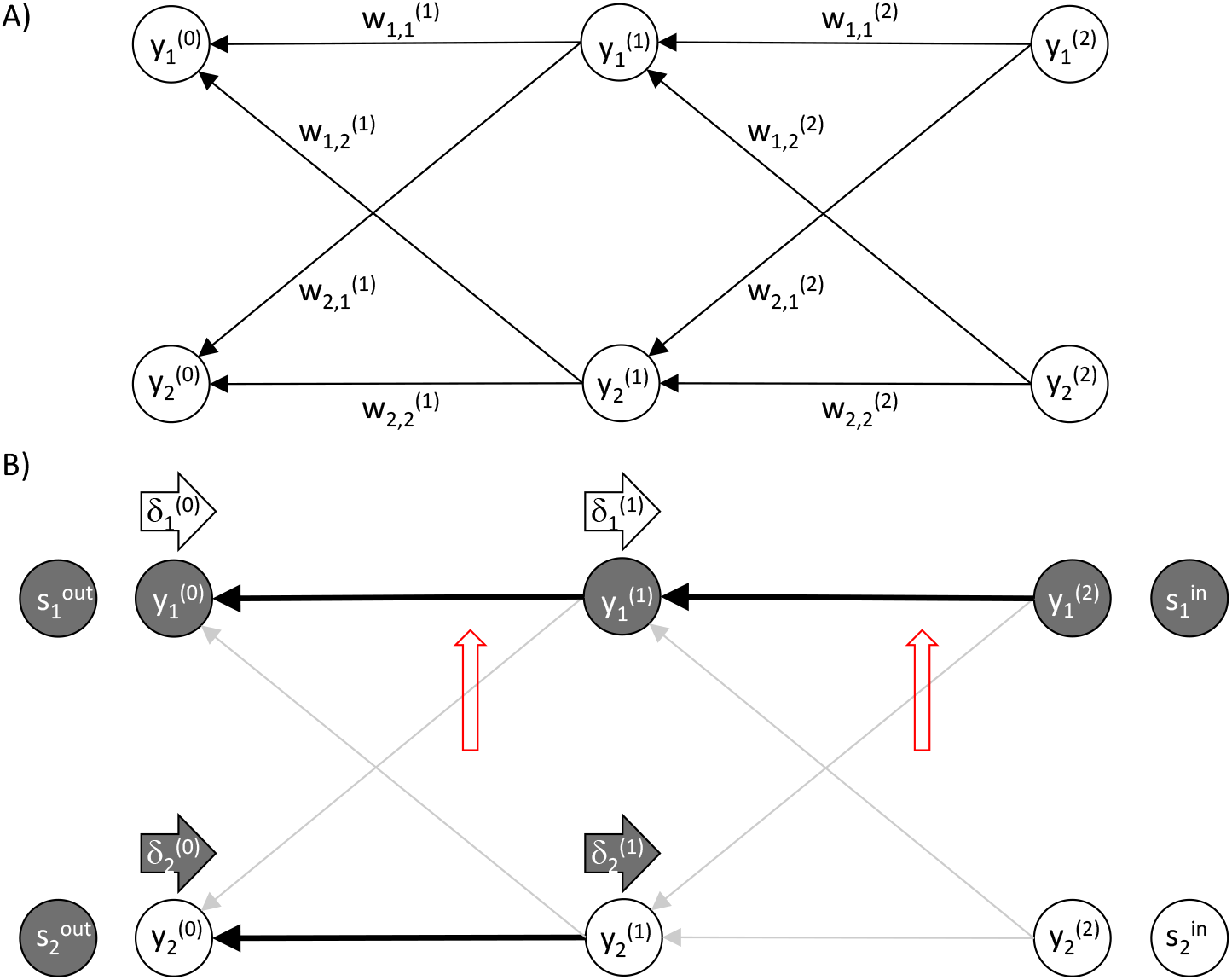
Back-propagation algorithm. A) Architecture of an ANN. Circles denote nodes and arrows denote connections. B) An example of activity and weight changes in an ANN. Thick black arrows between the nodes denote connections with high weights, while thin grey arrows denote the connections with low weights. Filled and open circles denote nodes with higher and lower activity, respectively. Rightward pointing arrows labelled 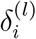 denote error terms and their darkness indicates how large the errors are. Upward pointing red arrows indicate the weights that would most increase according to the back-propagation algorithm.

where 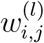 is the weight from the *j^th^* node in the *l^th^* layer to the *i^th^* node in the (*l* — 1)*^th^* layer, and *n*^(*l*)^ is the number of nodes in layer *l*. For brevity, we will refer to variable 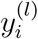, as the activity of a node.

The output the network produces for a given input depends on the values of the weight parameters. This can be illustrated in an example of an ANN shown in Figure 1B: The output node 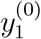 has a high activity as it receives an input from the active input node 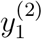 via strong connections. By contrast, for the other output node 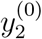 there is no path leading to it from the active input node via strong connections, so its activity is low.

The weight values are found during the following training procedure. At the start of each iteration, the activities in the input layer 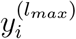 are set to values from input training sample, which we denote by 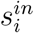. The network first makes a prediction, i.e. the activities of nodes are updated layer by layer according to Equation 1. The predicted output in the last layer 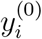 is then compared to the output training sample 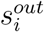 We wish to minimize the difference between the actual and desired output, so we define the following objective function: ^1^

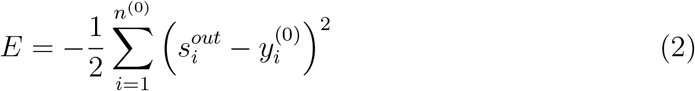

The training set contains many pairs of training vectors 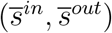, which are iteratively presented to the network, but for simplicity of notation we will consider just changes in weights after presentation of a single training pair. We wish to modify the weights to maximize the objective function, so we update the weights proportionally to the gradient of the objective function:

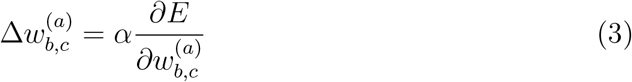

where *α* is a parameter describing the learning rate.

Since weight 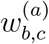 determines activity 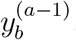, the derivative of the objective function over the weight can be found by applying the chain rule:

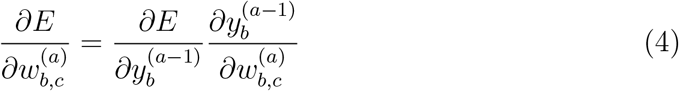

The first partial derivative on the right hand side of the above equation expresses by how much the objective function can be increased by increasing the activity of node *b* in layer *a* − 1, and we will denote it by:

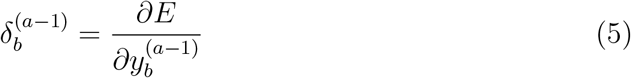

The values of these error terms for the sample network in Figure 1B are indicated by the darkness of the arrows labelled 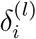. The error term 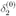 is high because there is a mismatch between the actual and desired network output, so by increasing the activity in the corresponding node 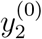 the objective function can be increased. By contrast the error term 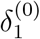 is low, because the corresponding node 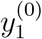 already produces the desired output, so changing its activity will not increase the objective function. The error term 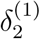 is high because the corresponding node 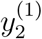 projects strongly to the node 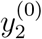 producing too low output, so increasing value of 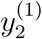 can increase the objective function. For analogous reasons, the error term 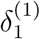 is low.

Now let us calculate the error terms 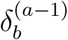. It is straightforward to evaluate them for the output layer:

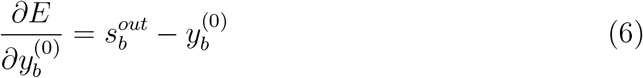

If we consider a node in an inner layer of the network then we must consider all possible routes through which the objective function is modified when the activity of the node changes, i.e. we must consider the total derivative

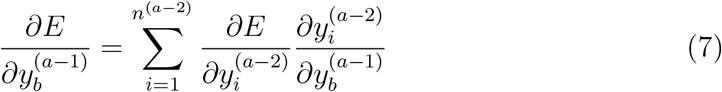

Using the definition of Equation (5), and evaluating the last derivative of the above equation using the chain rule, we obtain the recursive formula for the error terms:

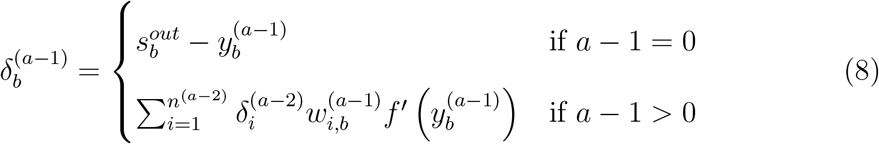

The fact that the error terms in layer *l* > 0 can be computed on the basis of the error terms in the next layer *l* − 1 gave the name: “error back-propagation” algorithm.

Substituting the definition of error terms from Equation 5 into Equation 4 and evaluating the second partial derivative on the right hand side of Equation 4 we obtain:

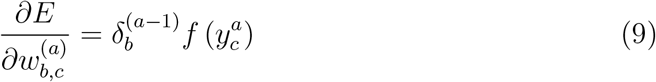

According to the above equation, the change in weight 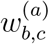 is proportional to the product of the output from the pre-synaptic node 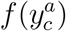, and the error term 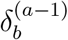 associated with the post-synaptic node. Red upward pointing arrows in Figure 1B indicate which weights would be most increased in this example, and it is evident that the increase in these weights will indeed increase the objective function.

In summary, after presenting to the network a training sample, each weight is modified proportionally to the gradient given in Equation 9 with the error term given by Equation 8. The expression for weight change (Equations 9 and 8) is a complex global function of activities and weights of neurons not connected to the synapse being modified. In order for real neurons to compute it, the architecture of the model could be extended to include nodes computing the error terms, that could affect the weight changes. As we will see, analogous nodes are present in the predictive coding model.

## Predictive coding for supervised learning

Due to the generality of the predictive coding framework, there are multiple network architectures within this framework that can perform supervised learning. In this section we describe the simplest model that can closely approximate the back-propagation, and we will consider other architectures later. The description in this section closely follows that of unsupervised predictive coding networks (Rao and Ballard, 1999; Friston, 2005), but is adapted for the supervised setting. Also, we provide a succinct description of the model, but for readers interested in a gradual and more detailed introduction to the predictive coding framework, we recommend reading sections 1-2 of a tutorial on this framework (Bogacz, 2017) before reading this section.

We first propose a probabilistic model for supervised learning, then we describe the inference in the model, its neural implementation, and finally learning of model parameters.

## Probabilistic model

Figure 2A shows a structure of a probabilistic model that parallels the architecture of the ANN shown in Figure 1A. It consists of *l_max_* layers of variables, such that the variables on level *l* depend on the variables on level *l* + 1. It is important to emphasize that Figure 2A does not show the architecture of the predictive coding network, but only the structure of underlying probabilistic model - as we will see below, the inference in this model can be implemented by a network with architecture shown in Figure 2B.

**Figure 2:**
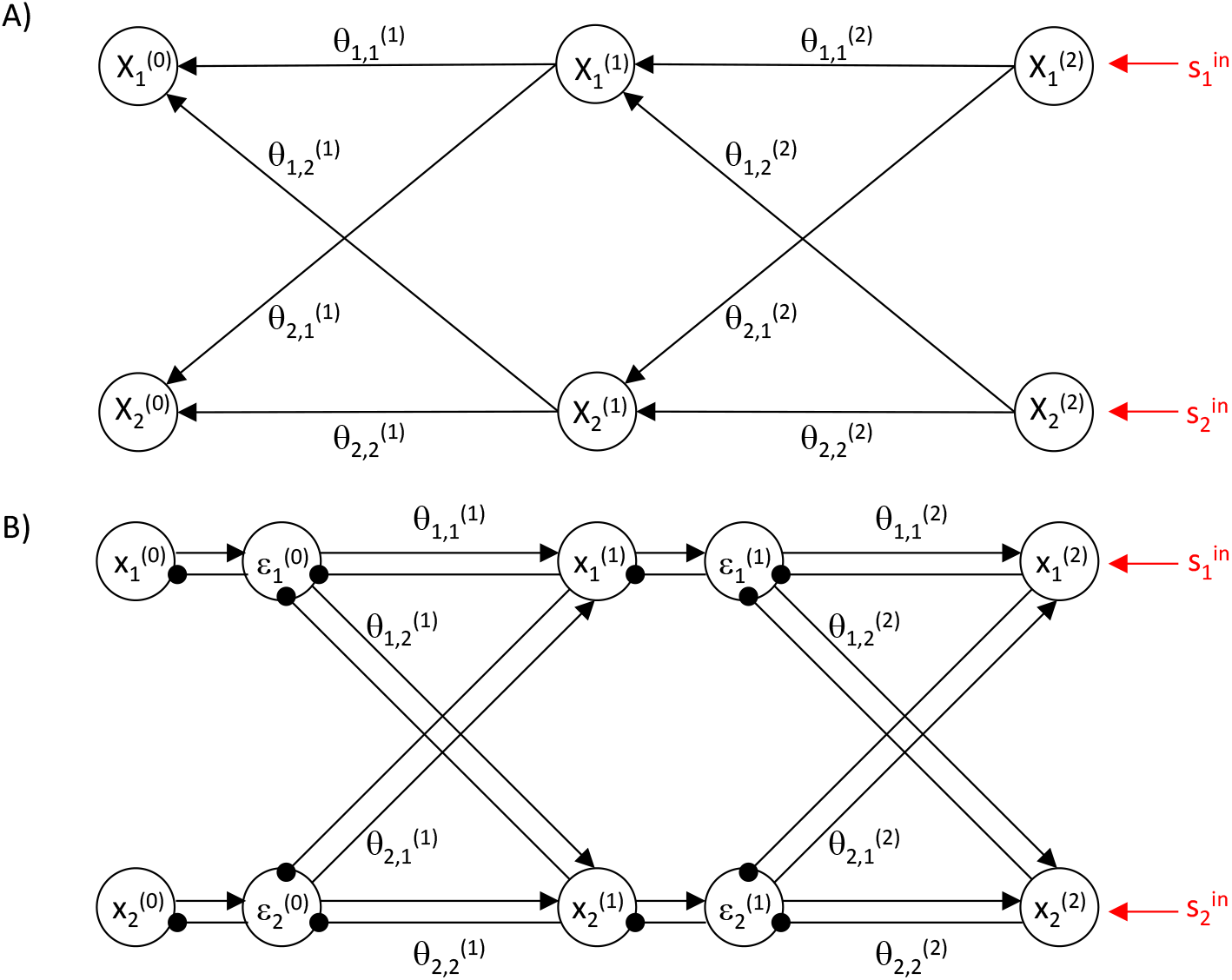
Predictive coding model. A) Structure of the probabilistic model. Circles denote random variables, while arrow denote dependencies between them. B) Architecture of the network. Arrows and lines ending with circles denote excitatory and inhibitory connections respectively. Connections without labels have weights fixed to 1.

By analogy to ANNs, we assume that variables on the highest level 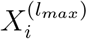 are fixed to the input sample 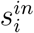, and the inferred values of variables on level 0 are the output from the network. Readers familiar with predictive coding models for sensory processing may be surprised that the sensory input is provided to the highest level, as traditionally in these models, the input is provided to level 0. Indeed, when biological neural networks learn in a supervised manner, both input and output are provided to sensory cortices. For example, when a child learns the sounds of the letters, the input (i.e. the shape of a letter) is provided to visual cortex, the output (i.e. the sound) is provided to the auditory cortex, and both of these sensory cortices communicate with associative cortex. The model we consider in this subsection corresponds to a part of this network: from associative areas to the sensory modality to which the output is provided. So in the example, level 0 corresponds to auditory cortex, while the highest levels correspond to associative areas. Thus the input 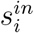 presented to this network does not correspond to raw sensory input, but rather to its representation pre-processed by visual networks. We will discuss how the sensory networks processing the input modality can be introduced to the model later in the Results section.

Let *^(l)^* be a vector of random variables on level *l*, and let us denote a sample from random variable *^(l)^* by *^(l)^*. Let us assume the following relationship between the random variables on adjacent levels (for brevity of notation we write *P*(*^(l)^*) instead of *P ^(l)^* = *^(l)^*):

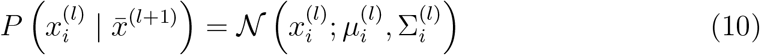

In the above equation *N*(*x; µ, Σ*) is the probability density of a normal distribution with mean *µ* and variance Σ. The mean of probability density on level *l* is a function of the values on the higher level analogous to the input to a node in ANN (cf. Equation 1):

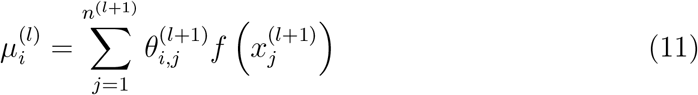

In the above equation *n*^(*l*)^ denotes the number of random variables on level *l*, and 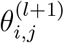 are the parameters describing the dependence of random variables. For simplicity in this paper we do not consider how 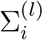 are learned (Friston, 2005; Bogacz, 2017), but treat them as fixed parameters.

### Inference

Let us now move to describing the inference in the model (i.e. finding most likely values of model variables, which will determine the activity of nodes in the predictive coding network). We wish to find the most likely values of all unconstrained random variables in the model, which maximize the probability *P*(*^(0)^*,…**^(*l_max_*–1)^|**^(*l_max_*)^). Since the nodes on the highest levels are fixed to 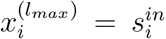, their values are not being changed but rather provide a condition on other variables. To simplify calculations we define the objective function equal to the logarithm of the joint distribution (since the logarithm is a monotonic operator, a logarithm of a function has the same maximum as the function itself):

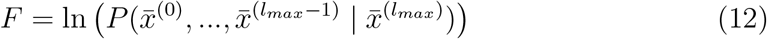

Since we assumed that the variables on one level just depend on variables of the level above, we can write the objective function as:

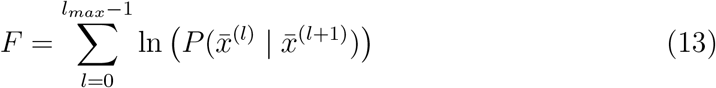

Substituting Equation 10 and the expression for a normal distribution into the above equation, we obtain:

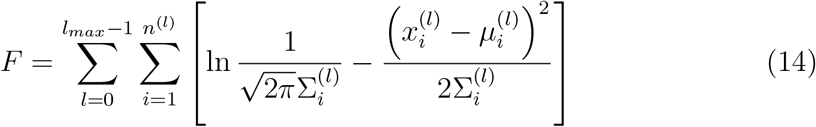

Then ignoring constant terms we can write the objective function as:

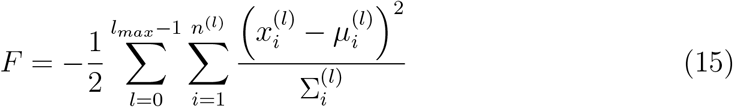

Recall that we wish to find the values 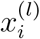 that maximize the above objective function. This can be achieved by modifying 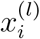 proportionally to the gradient of the objective function. To calculate the derivative of *F* over 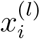 we note that each 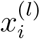 influences *F* in two ways: it occurs in Equation 15 explicitly, but it also determines the values of 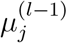. Thus the derivative contains two terms:

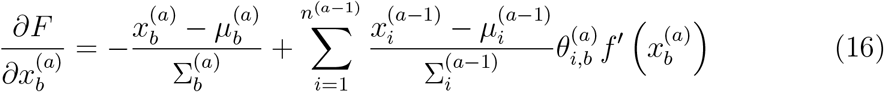

In the above equation, there are terms that repeat, so let us denote them by:

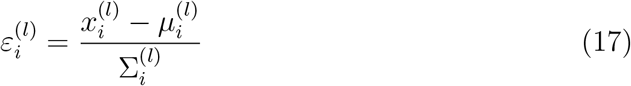

These terms describe by how much the value of a random variable on a given level differs from the mean predicted by a higher level, so let us refer to them as prediction errors. Substituting the definition of prediction errors into Equation 16 we obtain the following rule describing changes in 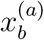 over time:

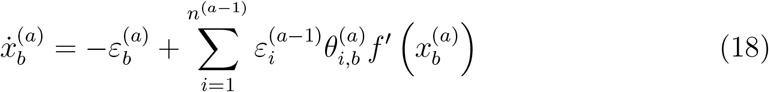

### Neural implementation

The computations described by Equations 17–18 could be performed by a simple network illustrated in Figure 2B with nodes corresponding to prediction errors 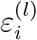 and values of random variables 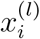. The prediction errors 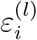 are computed on the basis of excitation from corresponding variable nodes 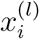, and inhibition from the nodes on the higher level 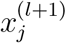 weighted by strength of synaptic connections 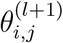 Conversely, the nodes 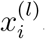 make computations on the basis of the prediction error from the corresponding level, and the prediction errors from the lower level weighted by synaptic weights.

It is important to emphasize that for a linear function *f*(*x*) = *x*, the non-linear terms in Equations 17–18 would disappear, and these equations could be fully implemented in the simple network shown in Figure 2B. To implement Equation 17, a prediction error node would get excitation from the corresponding variable node and inhibition equal to synaptic input from higher level nodes, thus it could compute the difference between them. Scaling the activity of nodes encoding prediction error by a constant 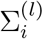 could be implemented by self-inhibitory connections with weight 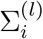 (we do not consider them here for simplicity - but for details see (Friston, 2005; Bogacz, 2017)). Analogously to implement Equation 18, a variable node would need to change its activity proportionally to its inputs.

One can imagine several ways how the non-linear terms can be implemented, and Figure 3 shows one of them (Bogacz, 2017). The prediction error nodes need to receive the input from the higher level nodes transformed through a non-linear function, and this transformation could be implemented by additional nodes (indicated by a hexagon labelled 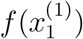 in Figure 3). Introduction of such additional nodes is also necessary to make the pattern of connectivity in the model more consistent with that observed in the cortex. In particular, in the original predictive coding architecture (Figure 2B), the projections from the higher levels are inhibitory, whereas connections between cortical areas are excitatory. Thus, to make the predictive coding network in accordance with this, the sign of the top down input needs to be inverted by local inhibitory neurons (Spratling, 2008). Here we propose that these local inhibitory neurons could additionally perform a non-linear transformation. With this arrangement, there are individual nodes encoding 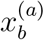 and 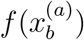 and each node only sends the value it encodes. According to Equation 18, the input from the lower level to a variable node needs to be scaled by a non-linear function of the activity of variable node itself. Such scaling could be implemented either by a separate node (indicated by a hexagon labelled 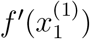 in Figure 3) or by intrinsic mechanisms within the variable node that would make it react to excitatory inputs differentially depending on its own activity level.

**Figure 3:**
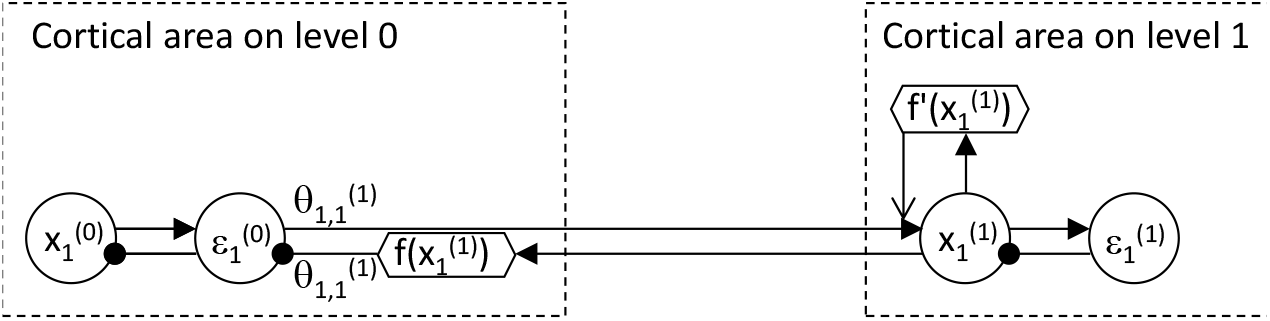
Possible implementation of non-linearities in the predictive coding model (magnification of a part of the network in Figure 2B). Filled arrows and lines ending with circles denote excitatory and inhibitory connections respectively. Open arrow denotes a modulatory connection with multiplicative effect. Circles and hexagons denote nodes performing linear and non-linear computations respectively.

In the predictive coding model, after the input is provided, all nodes are updated according to Equations 17–18, until the network converges to a steady state. We label variables in the steady state with an asterisk e.g. 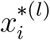 or *F*\*.

Figure 4A illustrates values to which a sample model converges when presented with a sample pattern. The activity in this case propagates from node 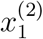 through the connections with high weights resulting in activation of nodes 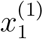 and 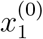 (note that double inhibitory connection from higher to lower levels has overall excitatory effect). Initially the prediction error nodes would change their activity, but eventually their activity converges to 0, as their excitatory input becomes exactly balanced by inhibition.

**Figure 4:**
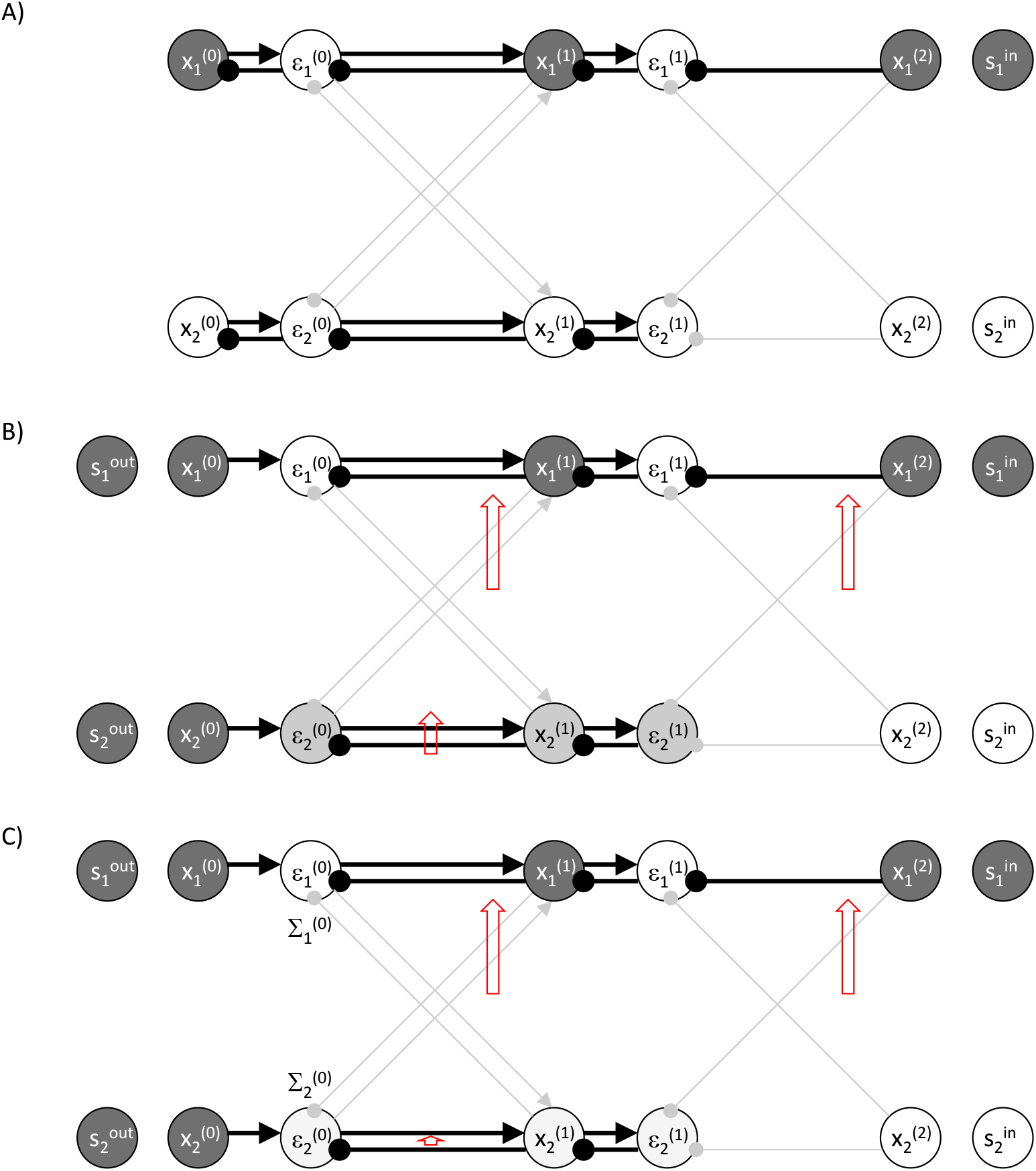
Example of a predictive coding network for supervised learning. A) Prediction mode. B) Learning mode. C) Learning mode for a network with high value of parameter describing sensory noise. Notation as in Figure 2B.

### Learning parameters

During learning, the values of the nodes on the lowest level are set to the output sample, i.e. 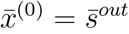, as illustrated in Figure 4B. Then the values of all nodes on levels *l* ∈{1, …, *l_max_* – 1 } are modified in the same way as described before (Equation 18).

Figure 4B illustrates an example of operation of the model. The model is presented with the desired output in which both nodes 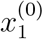 and 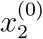 are active. Node 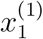 becomes active, as it receives both top down and bottom up input. There is no mismatch between these inputs, so the corresponding prediction error nodes (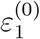 and 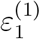) are not active. By contrast, the node 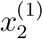 gets bottom up but no top down input, so its activity has intermediate value, and the prediction error nodes connected with it (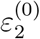 and 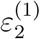) are active.

Once the network has reached its steady state, the parameters of the model 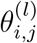 are updated so the model better predicts the desired output. This is achieved by modifying 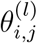 proportionally to the gradient of the objective function over the parameters. To compute the derivative of the objective function over 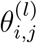, we note that 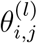 affects the value of function *F* of Equation 15 by influencing 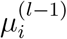, hence

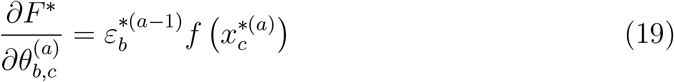

According to the above equation, the change in a synaptic weight 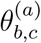 of connection between levels *a* and *a* − 1 is proportional to the product of quantities encoded on these levels. For a linear function *f* (*x*) = *x*, the non-linearity in the above equation would disappear, and the weight change would simply be equal to the product of the activities of pre-synaptic and post-synaptic nodes (Figure 2B). Even if the non-linearity is considered, as in Figure 3, the weight change is fully determined by the activity of pre-synaptic and post-synaptic nodes. The learning rules of the top and bottom weights must be slightly different. For the bottom connection labelled 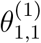 in Figure 3, the change in a synaptic weight is simply equal to the product of the activity of nodes it connects (round node 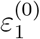 and hexagonal node 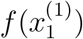). For the top connection, the change in weights is equal to the product of activity of the pre-synaptic node (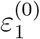) and function *f* of activity of the post-synaptic node (round node 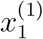). This then maintains the symmetry of the connections, i.e. the bottom and the top connections are modified by the same amount. We refer to these changes as Hebbian in a sense that in both cases the weight change is a product of monotonically increasing functions of activity of pre-synaptic and post-synaptic neurons.

Figure 4B illustrates resulting changes in the weights. In the example of Figure 4B, the weights that increase most are indicated by long red upward arrows. There would also be an increase in the weight between 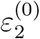 and 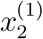, indicated by a shorter arrow, but it would be not as large as node 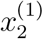 has lower activity. It is evident that after these weight changes the activity of prediction error nodes would be reduced indicating that the desired output is better predicted by the network. In Algorithm 1, we include pseudocode to clarify how the network operates in training mode.

#### Algorithm 1

Pseudocode for predictive coding during learning. Please note that in the simulations presented, to make for faster learning, first a prediction was made by inputting 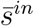 alone and propagating through the network layer by layer, as we know that all error nodes eventually would converge to zero in the prediction phase (see next section). Then the output 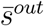 is applied, after which inference took place.

**for all** Data **do**

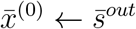

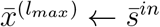

**repeat**

Inference: Equation 17, 18

**until** convergence

Update weights: Equation 19

## Results

### Relationship between the models

An ANN has two modes of operation: during prediction it computes its output on the basis of 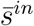, while during learning it updates its weights on the basis of 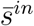 and 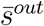. The predictive coding network can also operate in these modes. We now discuss the relationship between computations of an ANN and a predictive coding network in these two modes.

### Prediction

We now show that the predictive coding network has a stable fixed point at the state where all nodes have the same values as the corresponding nodes in the ANN receiving the same input 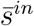. Since all nodes change proportionally to the gradient of *F*, the value of function *F* always increases. Since the network is constrained only by the input, the maximum value *F* can reach is 0, and because *F* is a negative of sum of squares, and this maximum is achieved if all terms in the summation of Equation 15 are equal to 0, i.e. when:

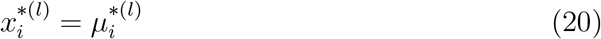

Since 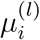 is defined in analogous way as 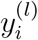 (cf. Equations 1 and 11), the nodes in the prediction mode have the same values at the fixed point as the corresponding nodes in the ANN, i.e. 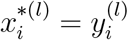.

The above property is illustrated in Figure 4A, in which weights are set to the same value as for the ANN in Figure 1B, and the network is presented with the same input sample. The network converges to the same pattern of activity on level *l* = 0 as for the ANN in Figure 1B.

### Learning

The pattern of weight change in the predictive coding network shown in Figure 4B is similar as in back-propagation algorithm (Figure 1B). Let us now analyse under what conditions weight changes in the predictive coding model converge to that in the back-propagation algorithm.

The weight update rules in the two models (Equations 9 and 19) have the same form, however, the prediction error terms 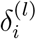 and 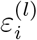 were defined differently. To see the relationship between these terms, we will now derive the recursive formula for prediction errors 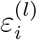 analogous to that for 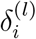 in Equation 8. We note that once the network reaches the steady state in the learning mode, the change in activity of each node must be equal to zero. Setting the left hand side of Equation 18 to 0 we obtain:

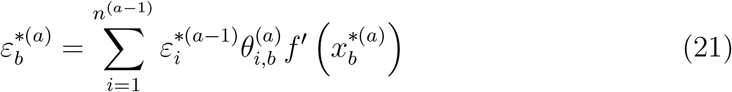

We can now write a recursive formula for the prediction errors:

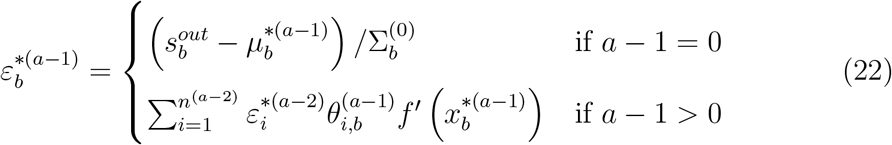

Let us first consider the case when all variance parameters are set to 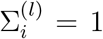 (as in (Rao and Ballard, 1999)). Then the above formula has exactly the same form as for the back-propagation algorithm (Equation 8). Therefore, it may seem that weight change in the two models is identical. However, for the weight change to be identical, the values of the corresponding nodes must be equal, i.e. 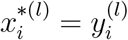 (it is sufficient for this condition to hold for *l* > 0, because 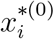 do not directly influence weight changes). Although we have shown in the previous subsection that 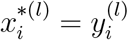 in the prediction mode, it may not be the case in the learning mode, because the nodes 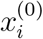 are fixed (to 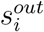), and thus function *F* may not reach the maximum of 0, so Equation 20 may not be satisfied.

Let us now consider under what conditions 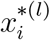 is equal or close to 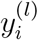. First, when the networks are trained such that they correctly predict the output training samples, then objective function *F* can reach 0 during the relaxation and hence 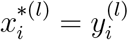, and the two models have exactly the same weight changes. In particular, the change in weights is then equal to 0, thus the weights resulting in perfect prediction are a fixed point for both models.

Second, when the networks are trained such that their predictions are close to the output training samples, then fixing 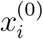 will only slightly change the activity of other nodes in the predictive coding model, so the weight change will be similar.

To illustrate this property we compare the weight changes in predictive coding models and ANN with very simple architecture shown in Figure 5A. This network consists of just three layers (*l_max_* = 2) and one node in each layer (*n*^(0)^ = *n*^(1)^ = *n*^(2)^ = 1). Such network has only 2 weight parameters (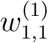 and 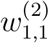), so the objective function of the ANN can be easily visualized. The network was trained on a set in which input training samples were generated randomly from uniform distribution 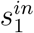 ε [−5, 5], and output training samples were generated as s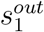 = *W*^(1)^ tanh(*W*^(2)^ tanh(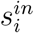), where *W*^(1)^ = *W*^(2)^ = 1 (Figure 5B). Figure 5C shows the objective function of the ANN for this training set. Thus an ANN with weights equal to 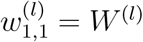 perfectly predicts all samples in the training set, so the objective function is equal to 0. There are also other combinations of weights resulting in good prediction, which create a “ridge” of the objective function.

**Figure 5:**
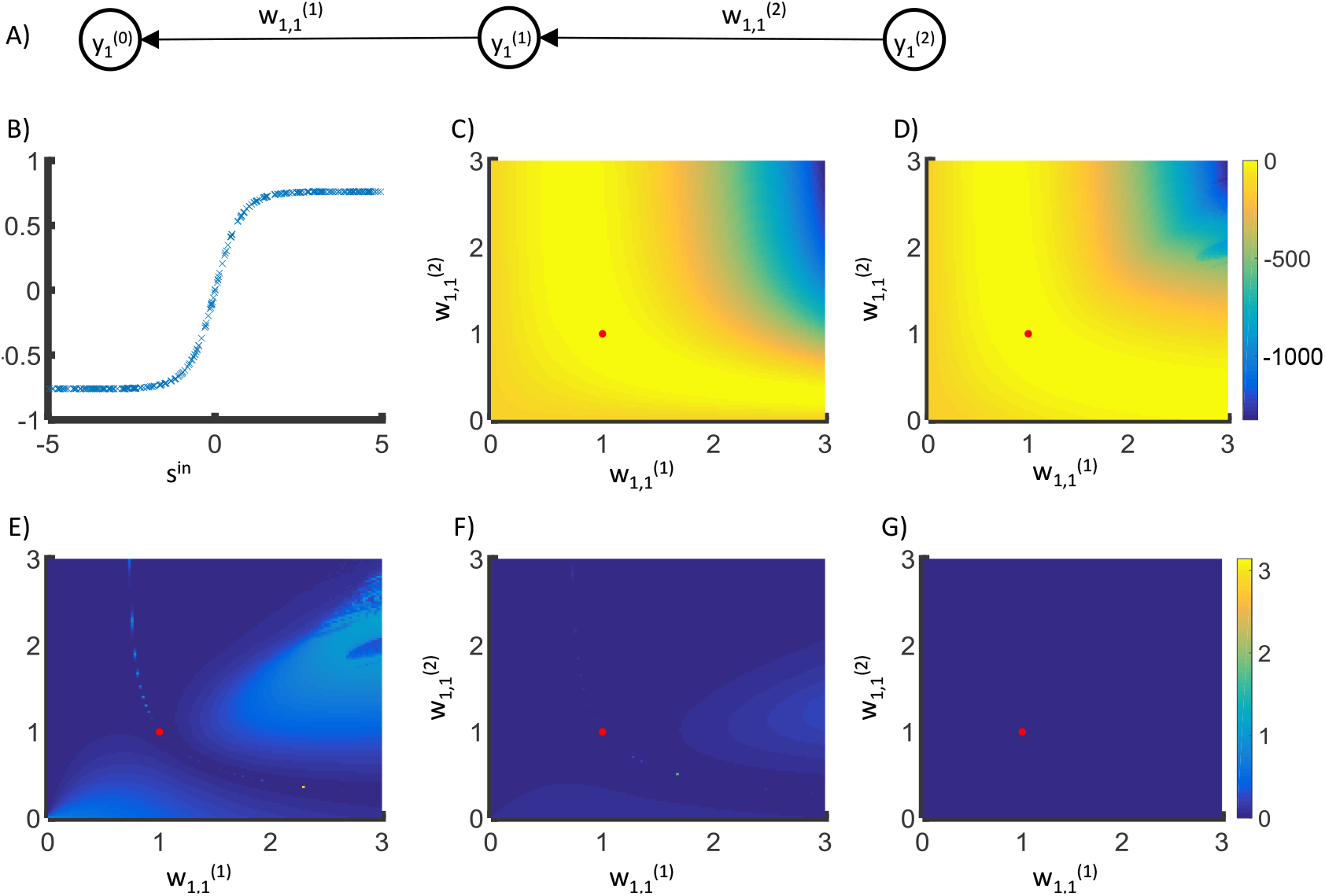
Comparison of weight changes in back-propagation and predictive coding models. A) The structure of the network used. B) The data that the models were trained on, here *s^out^* = *tanh*(*tanh*(*s^in^*)) C) The objective function of an ANN for a training set with 300 samples generated as described in main text. The objective function is equal to sum of 300 terms given by Equation 2 corresponding to individual training samples. The red dot indicates weights that maximize the objective function. D) The objective function of the predictive coding model at the fixed point. For each set of weights and training sample, to find the state of predictive coding network at the fixed point, the nodes in layers 0 and 2 were set to training examples, and the node in layer 1 was updated according to Equation 18. This equation was solved using Euler method. A dynamic form of the Euler integration step was used where its size was allowed to reduce by a factor should the system not be converging (i.e. the maximum change in node activity increases from the previous step). Initial step size was 0.2. The relaxation was performed until the maximum value of 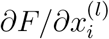 was lower than 10^−6^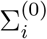 or 1,000,000 iterations had been performed. E-G) Angle difference between the gradient for the ANN and the gradient for the predictive coding model found from Equation 19. Different panels correspond to different values of parameter describing sensory noise: E) 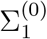 = 1. F) 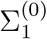 = 8. G) 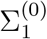 = 256.

Figure 5E shows the angle between the direction of weight change in back-propagation and the predictive coding model. The directions of the gradient for the two models are very similar except for the regions where the objective functions *E* and *F*\* are misaligned (cf. Figures 5C and D). Nevertheless close to the maximum of the objective function (indicated by a red dot), the directions of weight change become similar and the angle decreases towards 0.

There is also a third condition under which the predictive coding network approximates the back-propagation algorithm. Namely, when the value of parameters 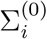 is increased relative to other 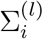, then the impact of fixing 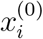 on the activity of other nodes is reduced, because 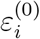 becomes smaller (Equation 17) and its influence on activity of other nodes is reduced. Thus 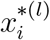 is closer to 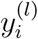 (for *l* > 0), and the weight change in the predictive coding model becomes closer to that in the back-propagation algorithm (recall that the weight changes are the same when 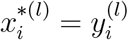 for *l* > 0).

Multiplying 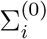 by a constant will also reduce all 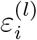 by the same constant (see Equation 22), and consequently all weight changes will be reduced by this constant. This can be compensated by multiplying the learning rate *α* by the same constant, so the magnitude of the weight change remains constant. In this case, the weight updates of the predictive coding network will become asymptotically similar to the ANN, regardless of prediction accuracy.

Figures 5F and G show that as 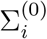 increases the angle between weight changes in the two models decreases towards 0. Thus as the parameters 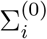 are increased, the weight changes in the predictive coding model converge to those in the back-propagation algorithm.

In the example of Figure 4, panel C illustrates the impact of increasing 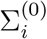. It reduces 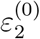, which in turn reduces 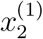 and 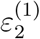. This decreases all weight changes, but particularly the change of the weight between nodes 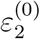 and 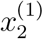 (indicated by a short red arrow) as both of these nodes have reduced activity. After compensating for the learning rate these weight changes become more similar to those in back-propagation algorithm (compare Figures 4B, C and 1B). We however note that learning driven by very small values of the error nodes is less biologically plausible. However in Figure 6 we will show that a high value of 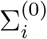 is not required for good learning with these networks.

**Figure 6:**
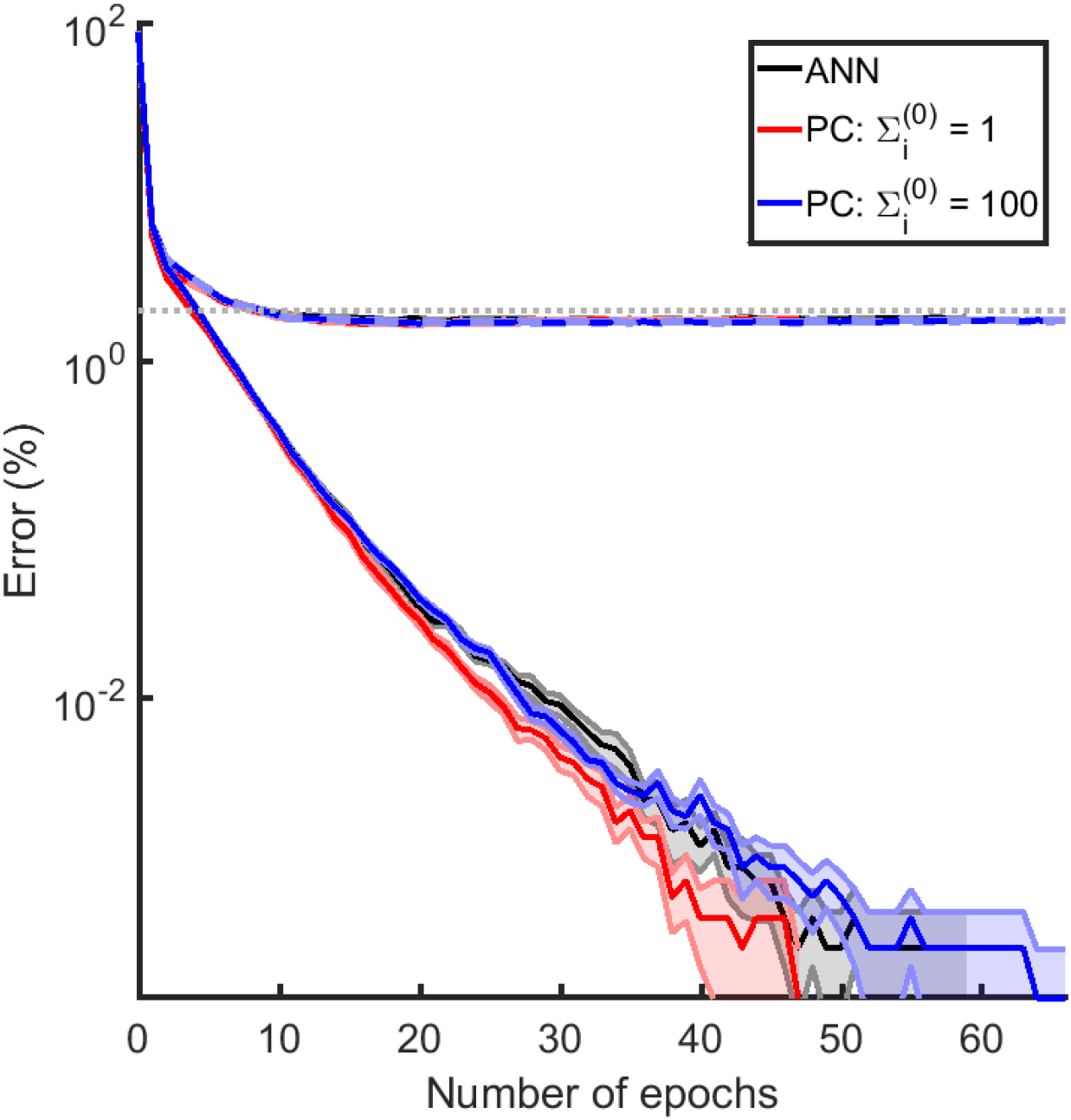
Comparison of prediction accuracy (%) for different models (indicated by colours - see key) on the MNIST dataset. Training errors are shown with solid lines, and validation errors with dashed lines. The dotted grey line denotes 2% error. The models were run 10 times each, initialised with different weights. When the training error lines stop, this is when the mean error of the 10 runs was equal to zero. The weights were drawn from a uniform distribution with maximum and minimum values of 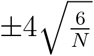 where *N* is the total number of neurons in the two layers either side of the weight. The input data was first transformed through an inverse logistic function as pre-processing, before being given to the network. When the network was trained with an image of class *c*, the nodes in layer 0 were set to: 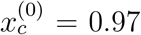 and 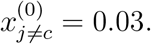. After inference and before the weight update, the error node values were scaled by 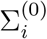 so as to be able to compare between the models. We used a batch size of 20, with a learning rate of 0.001 and the stochastic optimiser Adam (Kingma and Ba, 2014) to accelerate learning - this is essentially a per-parameter learning rate, where weights that are infrequently updated are updated more and vice-versa. We chose the number of steps in the inference phase to be 20, at this point the network will not necessarily have converged, but we did so to aid speed of training. This is not the minimum number of inference iterations that allows for good learning, this notion will be explored in a future paper. Otherwise simulations were as per Figure 5. The shaded regions in the fainter colour describe the standard error of the mean. The figure is shown on a logarithmic plot.

### Performance on more complex learning tasks

To efficiently learn in more complex tasks, ANNs include a “bias term” or an additional node in each layer which does not receive any input, but has activity equal to 1. Let us define this node as the node with index 0 in each layer, so 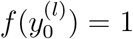. With such node, the definition of synaptic input (Equation 1) is extended to include one additional term 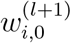, which is referred to as the “bias term”. The weight corresponding to the “bias term” is updated during learning according to the same rule as all other weights (Equation 9).

An equivalent “bias term” can be easily introduced to the predictive coding models. This would just be a node with a constant output of 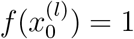 which projects to the next layer, but does have an associated error node. The activity of such node would not change after the training inputs are provided, and corresponding weights 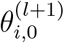 would be modified as all other weights (Equation 19).

To asses the performance of the predictive coding model on more complex learning tasks, we tested it on the MNIST dataset. This is a dataset of 28 by 28 images of handwritten digits, each associated with one of the 10 corresponding classes of digits. We performed the analysis for an ANN of size 784-600-600-10 (*l_max_* = 3), with predictive coding networks of the corresponding size too. We use the logistic sigmoid as the activation function. We ran the simulations for both the 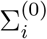 = 1 case and the 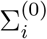 = 100 case. Figure 6 shows the learning curves for these different models. Each curve is the mean from ten simulations, with standard error shown as the shaded regions.

We see that the predictive coding models perform similarly to the ANN. For a large value of parameter 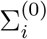 the performance of the predictive coding model was very similar to the back-propagation algorithm, in agreement with earlier analysis showing that then the weight changes in the predictive coding model converge to those in the back-propagation algorithm. Should we have had more than 20 steps in each inference stage, i.e. allowed the network to converge in inference, then the ANN and the predictive coding network with 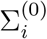 = 100 would have had an even more similar trajectory.

We see that all the networks eventually obtain a training error of 0.00%, and a validation error of ~ 1.7 – 1.8%. We did not optimise the learning rate for validation error as we are solely highlighting the similarity between ANNs and predictive coding.

### Effects of architecture of the predictive coding model

Since the networks we considered so far corresponded to the associative areas and sensory area to which the output sample was provided, the input samples 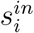 were provided to the nodes at the highest level of hierarchy, so we assumed that sensory inputs are already preprocessed by sensory areas. The sensory areas can be added to the model by considering an architecture in which there are two separate lower level areas receiving 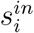 and 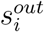, which are both connected with higher areas (de Sa and Ballard, 1998; Hyvarinen, 1999; O’Reilly and Munakata, 2000; Larochelle and Bengio, 2008; Bengio, 2014; Srivastava and Salakhutdinov, 2012; Hinton et al., 2006). For example, in case of learning associations between visual stimuli (e.g. shapes of letters) and auditory stimuli (e.g. their sounds), 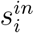 and 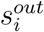 could be provided to primary visual and primary auditory cortices, respectively. Both of these primary areas project through a hierarchy of sensory areas to a common higher associative cortex.

To understand the potential benefit of such an architecture over the standard back-propagation, we analyse a simple example of learning the association between one dimensional samples shown in Figure 7A. Since there is a simple linear relationship (with noise) between samples in Figure 7A, we will consider predictions generated by a very simple network derived from a probabilistic model shown in Figure 7B. During training of this network the samples are provided to the nodes on the lowest level (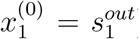 and 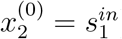).

**Figure 7:**
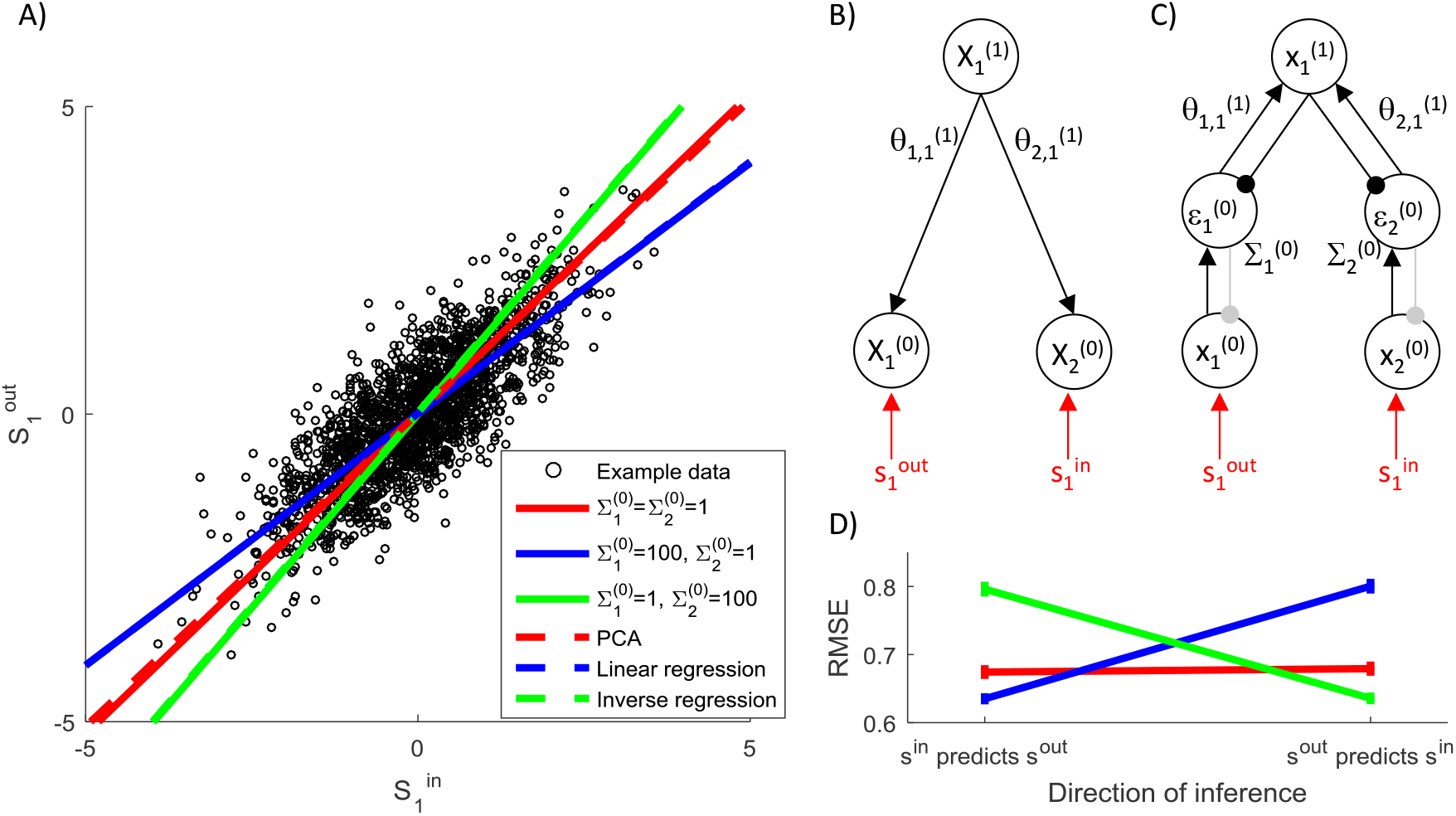
The effect of variance associated with different inputs on network predictions. A) Sample training set composed or 2000 randomly generated samples, such that 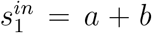 and 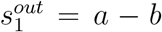 where *a* ˜ *N*(0,1) and *b* ˜ *N*(0,1/9). Lines compare the predictions made by the model with different parameters with predictions of standard algorithms (see key). B) Structure of probabilistic model and C) Architecture of the simulated predictive coding network. Notation as in Figure 2. Additionally, connections shown in grey are used if the network predicts the value of the corresponding sample. D) Root Mean Squared Error (RMSE) of the models with different parameters (see key of panel A) trained on data as in panel A and tested on further 100 samples generated from the same distribution. During the training, for each sample the network was allowed to converge to the fixed point as described in caption of Figure 5 and the weights were modified with learning rate *α* = 1. The entire training and testing procedure was repeated 50 times, and the error bars show the standard error.

For simplicity, we will assume a linear dependence of variables on the higher level:

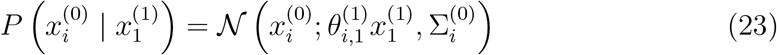

Since the node on the highest level is no longer constrained, we need to specify its prior probability, but for simplicity let us assume an uninformative flat prior 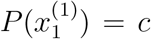, where *c* is a constant. Since the node on the highest level is uncon strained, the objective function we wish to maximize is the logarithm of the joint probability of all nodes:

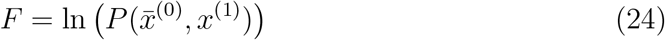

Ignoring constant terms this function has analogous form as in Equation 15:

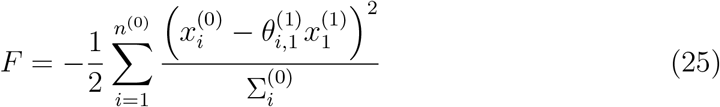

During training, the nodes on the lowest level are fixed, and node on the top level is updated proportionally to the derivative of *F*, analogously as in the models discussed previously:

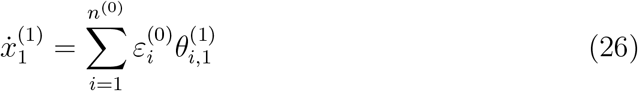

Analogously as before such computation can be implemented in a simple network shown in Figure 7C. After the nodes converge, the weights are modified to maximize *F*, which here is simply 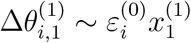.

During testing, we only set 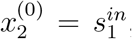, and let both nodes 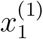 and 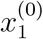 to be updated to maximize *F*, i.e. the node on the top level evolves according to Equation 26, while at the bottom level 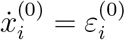.

Please note that this simple linear dependence could be captured by using a predictive coding network without a hidden layer and just by learning the means and covariance matrix i.e. 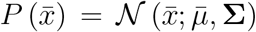, where 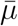 is the mean and Σ the covariance matrix. However we use a hidden layer to show the more general network, that could learn more complicated relationships if non-linear activation functions are used.

Solid lines in Figure 7A show the values predicted by the model (i.e. 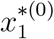) after providing different inputs (i.e 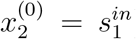),and different colours correspond to different noise parameters. When equal noise is assumed in input and output (red line), the network simply learns the probabilistic model that explains the most variance in the data, so the model learns the direction in which the data is most spread out. This direction is the same as the first principal component shown in dashed red line (any difference between the two lines is due the iterative nature of learning in the predictive coding model).

When the noise parameter at the node receiving output samples is large (blue line in Figure 7A), the dynamics of the network will lead to the node at the top level converging to the input sample (i.e. 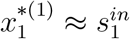). Given the analysis presented earlier, the model converges then to the back-propagation algorithm, which in the case of linear *f*(*x*) simply corresponds to linear regression, shown by dashed blue line.

Conversely, when the noise at the node receiving input samples is large (green line in Figure 7A), the dynamics of the network will lead to the node at the top level converging to the output sample (i.e. 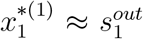). The network in this case will learn to predict the input sample on the basis of the output sample. Hence its predictions correspond to that obtained by finding linear regression in inverse direction (i.e. the linear regression predicting *s^in^* on the basis of *s^out^*), shown by the dashed green line.

Different predictions of the models with different noise parameters will lead to different amounts of error when tested, which are shown in the left part of Figure 7D (labelled *s^in^* predicts *s^out^*). The network approximating the back-propagation algorithm is most accurate, as the back-propagation algorithm explicitly minimizes the error in predicting output samples. Next in accuracy is the network with equal noise on both input and output, followed by the model approximating inverse regression.

Due to the flexible structure of the predictive coding network, we can also test how well it is able to infer the likely value of input sample *s^in^* on the basis of the output sample *s^out^*. In order to test it, we provide the trained network with the output sample (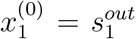), and let both nodes 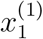 and 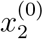 to be updated. The value 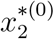 to which the node corresponding to the input converged is the network’s inferred value of the input. We compared these values with actual *s^in^* in the testing examples, and the resulting root mean squared errors are shown in the right part of Figure 7D (labelled *s^out^* predicts *s^in^*). This time the model approximating the inverse regression is most accurate.

Figure 7D illustrates that when noise is present in the data, there is a trade-off between accuracy of inference in the two directions. Nevertheless, the predictive coding network with equal noise parameters for inputs and outputs is predicting relatively well in both directions, being just slightly less accurate than the optimal algorithm for the given direction.

It is also important to emphasize that the models we analysed in this section generate different predictions, only because the training samples are noisy. If the amount of noise were reduced, the models’ predictions would become more and more similar (and their accuracy would increase). This parallels the property discussed earlier that the closer the predictive coding models predict all samples in the training set, the closer their computation to ANNs with back-propagation algorithm.

The networks in the cortex are likely to be non-linear and include multiple layers, but predictive coding models with corresponding architectures are still likely to retain the key properties outlined above. Namely, they would allow learning bidirectional associations between inputs and outputs, and if the mapping between the inputs and outputs could be perfectly represented by the model, the networks could be able to learn them and make accurate predictions.

## Discussion

In this paper we have proposed how the predictive coding models can be used for supervised learning. We showed that they perform the same computation as ANNs in the prediction mode, and weight modification in the learning mode has a similar form as for the back-propagation algorithm. Furthermore, in the limit of parameters describing the noise in the layer where output training samples are provided, the learning rule in the predictive coding model converges to that for the back-propagation algorithm.

### Biological plausibility of the predictive coding model

In this subsection we discuss various aspects of the predictive coding model that require consideration or future work to demonstrate the biological plausibility of the model.

In the first presented model (Subsection: Predictive coding for supervised learning) and in the simulations of hand-written digit recognition, the inputs and outputs corresponded to different layers to the traditional predictive coding model (Rao and Ballard, 1999), where the sensory inputs are presented to layer l = 0 while the higher layers extract underlying features. However, as mentioned while introducing the model, supervised learning in a biological context would often involve presenting the stimuli to be associated (e.g. image of a letter, and a sound) to sensory neurons in different modalities, and thus would involve the network from “input modality” via the higher associative cortex to the “output modality”. We focussed in this paper on analysing a part of this network from the higher associative cortex to the “output modality”, and thus we presented *s^out^* to nodes at layer *l* = 0. We did this because only for this case it is easy to show analytically the relationship between predictive coding and ANNs. Nevertheless we would expect the predictive coding network to also perform supervised learning when *s^in^* is presented to layer 0, while *s^out^* to layer *l_max_*, because the model minimizes the errors between predictions of adjacent levels so it learns the relationships between the variables on adjacent levels. It would be an interesting direction for a future work to compare the performance of the predictive coding networks with input and outputs presented to different layers.

In the last subsection of the results we briefly considered a more realistic architecture involving both modalities represented on lowest level layers. Such an architecture would allow for a combination of supervised and unsupervised learning. If one no longer has a flat prior on the hidden node, but a Gaussian prior (so as to specify a generative model), then each arm could be trained separately in an unsupervised manner, while the whole network could also be trained together. Consider now that the input to one of the arms is an image, and the input at the other arm is the classification. It would be interesting to investigate if the image arm could be pre-trained separately in an unsupervised manner alone, and if this would speed up learning of the classification.

Let us now consider the model in the context of the plausibility criteria stated in the Introduction. The first two criteria of local computation and plasticity are naturally satisfied in a linear version of the model (with *f*(*x*) = *x*), and we discussed possible neural implementation of non-linearities in the model (Figure 3). In that implementation some of the neurons have a linear activation curve (like the value node 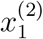 in Figure 3) and others are non-linear (like the node *f*(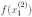)), which is consistent with the variability of firing-Input relationship (or f-I curve) observed in biological neurons (Bogacz et al., 2016).

The third criterion of minimal external control is also satisfied by the model, as it performs computations autonomously given input and outputs. The model can also autonomously “recognize” when the weights should be updated, because this should happen once the nodes converged to an equilibrium and have stable activity. It is interesting to point out that this simple rule would result in weight update in the learning mode, but no weight change in the prediction mode, because then the prediction error nodes have activity equal to 0, so the weight change (Equation 19) is also 0. Nevertheless, without a global control signal, each synapse could only detect if the two neurons it connects have converged. It will be important to investigate if such a local decision of convergence is sufficient for good learning.

The fourth criterion of plausible architecture is more challenging for the predictive coding model. First, the model includes special one-to-one connections between variable nodes (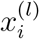) and the corresponding prediction error nodes (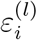), while there is no evidence for such special pairing of neurons in the cortex. It would be interesting to investigate if the predictive coding model would still work if these one-to-one connections were replaced by distributed ones. Second, the mathematical formulation of the predictive coding model requires symmetric weights in the recurrent network, while there is no evidence for such a strong symmetry in cortex. However, our preliminary simulations suggest that symmetric weights are not necessary for good performance of predictive coding network (as we will discuss in a forthcoming paper). Third, the error nodes can be either positive or negative, while biological neurons cannot have negative activity. Since the error neurons are linear neurons,and we know that rectified linear neurons exist in biology (Bogacz et al., 2016), we can approximate a purely linear neuron in the model with a biological rectified linear neuron if we associate zero activity in the model with baseline firing rate of a biological neuron. It will be important to test if such an approximation still results in efficient computation.

Nevertheless, predictive coding is an appealing framework for modelling cortical networks, as it naturally describes hierarchical organisation consistent with those of cortical areas (Friston, 2003). Furthermore, responses of some cortical neurons resemble those of prediction error nodes, as they show a decrease in response to repeated stimuli (Brown and Aggleton, 2001; Miller and Desimone, 1993), and increase in activity to unlikely stimuli (Bell et al., 2016). Additionally, neurons have been recently reported in the primary visual cortex which respond to a mismatch between actual and predicted visual input (Fiser et al., 2016; Zmarz and Keller, 2016).

### Does the brain implement back-prop?

This paper shows that a predictive coding network converges to back-propagation in a certain limit of parameters. However, it is important to add that this convergence in the limit is more of a theoretic result, as for parameters it occurs, the activity of error nodes becomes close to 0, so it is unclear if real neurons encoding information is spikes could reliably encode the prediction error. Nevertheless, the conditions under which the predictive coding model converges to the back-propagation algorithm are theoretically useful, as they provide an alternate probabilistic interpretations of the back-propagation algorithm. This allows both a comparison of the assumptions made by the back-propagation algorithm with the probabilistic structure of learning tasks, and questions whether setting the parameters of the predictive coding models to those approximating back-propagation is the most suitable choice for solving real-world problems faced by animals.

First, the predictive coding model corresponding to back-propagation assumes that output samples are generated from a probabilistic model with multiple layers of random variables, but most of the noise is added only at the level of output samples (i.e. 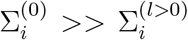). By contrast, probabilistic models corresponding to most of real-world datasets have variability entering on multiple levels. For example, if we consider classification of images of letters, the variability is present in both high level features like length or angle of individual strokes, and low level features like the colors of pixels.

Second, the predictive coding model corresponding to back-propagation assumes layered structure of the probabilistic model. By contrast probabilistic models corresponding to many problems may have other structures. For example, in the task from the Introduction of a child learning the sounds of the letters, the noise or variability is present in both the visual and auditory stimuli. Thus this task could be described by a probabilistic model including a higher level variable corresponding to a letter, which determines both the mean visual input perceived by a child, and the sound made by the parent. Thus the predictive coding networks with parameters that do not implement back-propagation algorithm exactly may be more suited for solving the learning tasks faced by animals and humans.

In summary, the above analysis suggests that it is unlikely that brain networks implement the back-propagation algorithm exactly. Instead, it seems more probable that cortical networks perform computations similar to those of a predictive coding network without any variance parameters dominating any others. These networks would be able to learn relationships between modalities in both directions, and flexibly learn probabilistic models well describing observed stimuli and the associations between them.

### Previous work on approximation of the back-propagation algorithm

As mentioned in the Introduction, other models have been developed describing how the back-propagation algorithm could be approximated in a biological neural network. We now review these models, relate them to the four criteria stated in the Introduction, and compare them with the predictive coding model.

O’Reilly (1998) considered a modified ANN which additionally includes feedback weights between layers which are equal to feed-forward weights. In this modified ANN, the output of hidden nodes in the equilibrium is given by

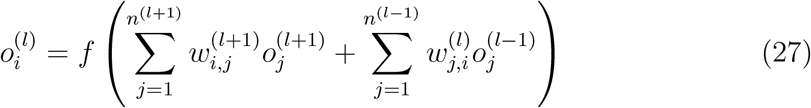

and the output of the output nodes satisfies in equilibrium the same condition as for the standard ANN (an equation similar to the one above but just including the first summation). It has been demonstrated that the weight change minimizing the error of this network can be well approximated by the following update (O’Reilly, 1998):

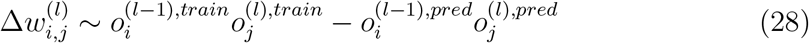

This is the contrastive Hebbian learning weight update rule (Ackley et al., 1985). In the above equation, 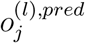 denotes the output of the nodes in the prediction phase, when the input nodes are set to 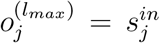 and all the other nodes are updated as described above, while 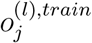 denotes the output in the training phase when additionally the output nodes are set to 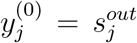, and the hidden nodes satisfy Equation 27. Thus according to the above plasticity rule, each synapse needs to be updated twice, once after the network settles to equilibrium during prediction, and once after the network settles following the presentation of the desired output sample. Each of these two updates relies just on local plasticity, but they have the opposite sign. Thus the synapses on all levels of hierarchy need “to be aware” of the presence of *s^out^* on the output, and use Hebbian or anti-Hebbian plasticity accordingly. Although it has been proposed how such plasticity could be implemented (O’Reilly, 1998), it is not known if cortical synapses can perform such form of plasticity.

In the above GeneRec model the error terms δ are not explicitly represented in neural activity, and instead the weight change based on errors is decomposed into a difference of two weight modifications: one based on target value and one based on predicted value. By contrast, the predictive coding model includes additional nodes explicitly representing error, and thanks to them has a simpler plasticity rule involving just a single Hebbian modification. A potential advantage of such single modification is robustness to uncertainty about the presence of *s^out^* as no mistaken weight updates can be made when *s^out^* is not present.

Bengio and colleagues (Bengio, 2014; Bengio et al., 2015) considered how the back-propagation algorithm can be approximated in a hierarchical network of auto-encoders, which learn to predict their own inputs. The general frameworks of auto-encoders and predictive coding are closely related, as both of the networks, which include feed-forward and feedback connections, learn to predict activity on lower levels from the representation on the higher levels. This work (Bengio, 2014; Bengio et al., 2015) includes many interesting results such as improvement of learning due to addition of noise to the system. However, it was not described how it is mapped on a network of simple nodes performing local computation. There is a discussion of a possible plasticity rule at the end of (Bengio, 2014), which has a similar form as Equation 28 of the GeneRec model.

Bengio and colleagues (Scellier and Bengio, 2016; Bengio and Fischer, 2015) introduce another interesting approximation to implement back-propagation in biological neural networks. It has some similarities to the model presented here in that it minimises an energy function. It however, like contrastive Hebbian learning, operates in two phases - a positive and a negative phase, where weights are updated from information obtained from each phase. The weights are changed following a differential equation update starting at the end of the negative phase and until convergence of the positive phase. Learning must be inhibited during the negative phase, which would require a global signal. This model also achieves good results on the MNIST dataset.

Lillicrap et al. (2014) focussed on addressing the requirement of the back-propagation algorithm that the error terms need to be transmitted backwards through exactly the same weights that are used to transmit information feed-forward. Remarkably, they have shown that even if random weights are used to transmit the errors backward, the model can still learn efficiently. Their model requires external control over nodes to route information differentially during training and testing, so it does not satisfy the third criterion stated in the Introduction. Furthermore, we note that the requirement of symmetric weights between the layers can be enforced by using symmetric learning rules like those proposed in GeneRec and predictive coding models. Equally, we will show in a future paper that the symmetric requirement is not actually necessary in the predictive coding model.

Balduzzi et al. (2014) showed that efficient learning may be achieved by a network which receives a global error signal, and in which synaptic weight modification depends jointly on the error and the terms describing the influence of each neuron of final error. However, it is not specified in this paper how these influence terms could be computed in a way satisfying the criteria stated in the Introduction.

Finally, it is worth pointing out that previous papers have shown that certain models perform similar computations as ANNs or that they approximate the back-propagation algorithm, while here we for a first time we show that a biologically plausible algorithm may actually converge to back-propagation. Although, this convergence in the limit is more of a theoretic result, it provides a mean to clarify the computational relationship between the proposed model and back-propagation, as described above.

### Relationship to experimental data

We hope that the proposed extension of the predictive coding framework to supervised learning will make it easier to experimentally test this framework. The model predicts that in a supervised learning task, like learning sounds associated with shapes, the activity after feedback, proportional to the error made by a participant, should be seen not only in auditory areas but also visual and associative areas. In such experiments, the model can be used to estimate prediction errors, and one could analyse precisely which cortical regions or layers have activity correlated with model variables. Inspection of the neural activity could in turn refine the predictive coding models, so they better reflect information processing in cortical circuits.

The proposed predictive coding models are still quite abstract and it is important to investigate if different linear or non-linear nodes can be mapped on particular anatomically defined neurons within a cortical micro-circuit (Bastos et al., 2012). Iterative refinements of such mapping on the basis of experimental data (such as f-I curves of these neurons, their connectivity and activity during learning tasks) may help understand how supervised and unsupervised learning is implemented in the cortex.

Predictive coding has been proposed as a general framework for describing computations in the neocortex (Friston, 2010). It has been shown in the past how networks in the predictive coding framework can perform unsupervised learning, attentional modulations, and action selection (Rao and Ballard, 1999; Feldman and Friston, 2010; Friston et al., 2010). Here we add to this list supervised learning, and associative memory (as the networks presented here are able to associate patterns of neural activity with each other). It is remarkable that the same basic network structure can perform this variety of the computational tasks, also performed by the neocortex. Furthermore, this network structure can be optimized for different tasks by modifying proportions of synapses among different neurons. For example, the networks considered here for supervised learning did not include connections encoding covariance of random variables, which are useful for certain unsupervised learning tasks (Bogacz, 2017). These properties of the predictive coding networks parallel organization of the neocortex, where the same cortical structure is present in all cortical areas, only differing in proportions and properties of neurons and synapses in different layers.

## Acknowledgements

This work was supported by Medical Research Council grant MC UU 12024/5 and the EPSRC. The authors thank Tim Vogels, Chris Summerfield and Eduardo Martin Moraud for reading the previous version of the manuscript and for providing very useful comments.

## Footnotes

1 As in previous work linking the back-propagation algorithm to probabilistic inference (Rumelhart et al., 1995), we consider the output from the network to be 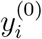 rather than *f*(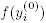), as it simplifies the notation of the equivalent probabilistic model. This corresponds to an architecture in which the nodes in the output layer are linear. A predictive coding network approximating an ANN with non-linear nodes in all layers was derived in a previous version of this paper (Whittington and Bogacz, 2015).

